# A Parametric Robustness versus Dynamic Sensitivity Paradox in a Bistable Biomolecular Circuit

**DOI:** 10.1101/2025.01.21.634208

**Authors:** Gunjan Chorasiya, Rudra Prakash, Shaunak Sen

## Abstract

Achieving robustness to multi-parametric perturbations where all parameters can change at the same time is challenging because the controller would also face the same disturbance as the plant. For nonlinear positive feedback, an important mechanism for cell fate determination in biomolecular contexts, quantitative aspects of robustness to such perturbations are generally unclear. Here we used mathematical methods of control and dynamical systems, interval analysis, and a benchmark model of a bistable biomolecular positive feedback circuit to address this. We confirmed that such perturbations can change the qualitative behaviour of the system extinguishing bistability. We obtained a quantitative relation between the relative variation in the stable steady state and the unstable steady state in terms of the relative changes in the parameters. We showed how the deviation in the trajectories near the unstable steady state due to multi-parametric perturbations could diverge almost exponentially after an initial transient, which could have a significant impact on the bistable switching dynamics. We found that the size of the eigenvalue for the unstable steady state was greater than that for the stable steady state, and proved this for certain parameters using a rigorous numerical construction. We noted a tradeoff between enhancing the parameter space of bistability and the increased sensitivity in the bistable dynamics due to parametric perturbations. We obtained rigorous bounds on the entire transient response for multi-parametric perturbations. These results provide a quantitative insight into the robustness of a bistable biomolecular positive feedback circuit to multi-parametric perturbations.

## I. Introduction

By making reversible processes irreversible and introducing the notion of memory, the idea of two stable steady states, or bistability, has provided a useful metaphor for investigations in multiple biological contexts, especially in the determination of cell fates and in signaling (Ferrell and Xiong, 2001). Bistability has been experimentally observed (Xiong and Ferrell, 2003; Ozbudak et al., 2004) and designed (Gardner et al., 2000; Zhu et al., 2022) in many different types of cells. From a dynamical systems point of view, bistability is typically associated with a saddle-node bifurcation (Strogatz, 2024; Ferrell, 2012). Optimal experimental design techniques have been used to detect bistability and provide a strategy to avoid uncertainty caused by bifurcations (Braniff et al., 2019). The key systems-level design principle is to have a positive feedback with sufficient nonlinearity in the underlying biomolecular circuit for bistability to be observed (Thomas and D’Ari, 2024; Alon, 2006). The nonlinear positive feedback serves to amplify specific input signals and triggers the transition from one state to another. This amplified sensitivity might also make the system sensitive to perturbations, both specific perturbations and multi-parametric perturbations, such as due to temperature where all parameters might change simultaneously.

Significant advances have been made in obtaining theoretical conditions for the existence of bistability in the nonlinear mathematical models of biomolecular circuits without having to explicitly solve them. The theory of monotone dynamical systems exploits the monotonic nature of interactions in biomolecular systems to obtain conditions for the existence of bistability (Angeli et al., 2004; Angeli and Sontag, 2004). Tools of algebraic geometry such as Sturm’s theorem have also been used to obtain conditions for the existence of bistability (Siegal-Gaskins et al., 2015). Recent work has also used the Jacobian determinant to derive algebraic conditions guaranteeing the existence of number of equilibrium points under state-dependent perturbations (McBride and Del Vecchio, 2020). These theoretical results have also been supplemented by extensive numerical studies, both in specific biomolecular contexts (Arkin et al., 1998) as well as in highlighting the role of various dynamical phenomena such as unstable saddle points in the dynamics (Trotta et al., 2012). (Proverbio et al., 2024) have performed the quantitative probabilistic assessment of the dynamical system’s ability to preserve a prescribed attractor under stochastic perturbations and introduced the resilience definition that generalises the notion of probabilistic robustness under stochastic perturbations effectively applied in biological models for noisy dynamics. These results provide a framework to understand and design bistability using biomolecular positive feedback circuits.

There are three striking aspects of bistable dynamics in biomolecular positive feedback circuits. One is the pervasiveness of these dynamics in multiple natural contexts and, increasingly, in synthetically designed contexts as well, where they may experience multi-parametric perturbations. Two is the impressive body of results that seeks to obtain conditions for the existence of bistability without needing to explicitly analytically solve the nonlinear equations in the mathematical models. Three is the importance of the unstable steady state, which while cannot be observed makes its influence on the bistable dynamics keenly felt. The quantitative robustness of these features to multi-parametric perturbations is generally unclear.

Here, we quantitatively investigated how design choices govern the sensitivity of bistable switching dynamics to multi-parametric perturbations. We addressed this using a benchmark nonlinear positive feedback model exhibiting bistable dynamics. Using the method of root locus, we obtained a necessary and sufficient condition for bistability based on the system parameters. We obtained a quantitative relation between the relative changes in the stable steady state and the unstable steady state in terms of the relative changes in the parameters. We showed that the deviation in the trajectories near the unstable steady state due to parametric perturbations increased almost exponentially, corresponding to the eigenvalue, but eventually bounded by the nonlinear dynamics. We also showed that the influence of divergence near an unstable state is reflected in the actual dynamics, exhibiting a significant difference between the perturbed and the unperturbed trajectories. The deviations in the transient dynamics near the unstable state can also be illustrated with stochastic simulations performed by Gillespie’s algorithm. We used interval analysis to obtain rigorous bounds on steady-state deviations under simultaneous multi-parameter variations. Across all Hill coefficients considered, the unstable eigenvalue consistently had a larger magnitude than the stable one, revealing an inherent trade-off between bistability robustness and switching sensitivity. We also used Taylor-model-based reachability analysis to enclose the system’s time trajectories under multi-parametric uncertainty. These results were illustrated using numerical simulations. We discussed the significance of the sensitivity of bistable dynamics to perturbations as well as a tradeoff between the size of the bistable regime in the parameter space and the sensitivity of the bistable dynamics.

## II. THEORY

Mathematical models of biomolecular circuits exhibiting bistable dynamics can have many state variables (for example, see (Siegal-Gaskins et al., 2015),(McBride and Del Vecchio, 2020; Arkin et al., 1998)). The typical bifurcation underlying these instances of bistability, however, are saddle-node bifurcations, which are essentially one dimensional (Strogatz, 2024).

### A. Existence of Bistability

For the above reason and for simplicity, we consider and analyse a one dimensional model of bistable dynamics, which is widely considered to be a benchmark model (Thomas and D’Ari, 2024; Alon, 2006). The differential equation in this mathematical model is

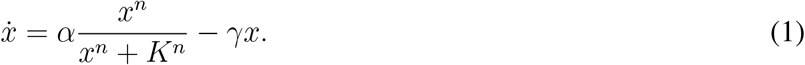

Here, *x* is the concentration of the protein that can activate its own expression. This is modelled with a Hill-type production function with co-operativity *n*, maximum production rate *α*, and DNA binding constant *K*. The rate of change of *x* is the difference between its production rate and degradation rate. The degradation rate is modelled as a first order process with rate constant *γ*. Because the production rate is an increasing function of *x*, an increase in *x* has a positive effect on the rate of change of *x*, implementing the positive feedback.

A qualitative geometric analysis can be used to show that such a system can exhibit bistability if the positive feedback has sufficient nonlinearity. In the bistable regime, there are three steady states *x*_*o*_, *x*_*u*_, and *x*_*s*_, corresponding to the points of intersection between the production rate and degradation rate (Alon, 2006). As this is a one dimensional system, their stabilities can be inferred from the slopes at the steady states and how they impact the rate of change of *x* in the neighbourhood of the steady states. As the slopes at the steady states *x*_*o*_ and *x*_*s*_ (denoted *λ*_*o*_ and *λ*_*s*_, respectively) are negative, they are stable steady states. The slope at *x*_*u*_ (denoted *λ*_*u*_) is positive and so this is an unstable steady state. These slopes explicitly appear in terms of the system parameters in the linearization of the dynamics in (1) around the steady states,

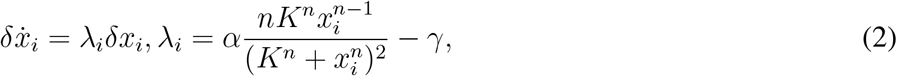

where *x*_*i*_, *i* ∈ {*o, s, u*} is the steady state and *δx*_*i*_ = *x* − *x*_*i*_ is the deviation around the steady state. From a functional standpoint, *x*_*o*_ and *x*_*s*_ represent two different cellular states, and *x*_*u*_ represents the threshold that needs to be crossed to transit from one cellular state to another. We presented an alternative proof of how the existence of bistability depends on system parameters. Our approach is based on a simple root locus construction to derive a proposition that explicitly characterizes the bistable regimes without the need for computing the steady-state solutions.

#### Proposition 1.

*The necessary and sufficient condition for bistability in* (1) *is* 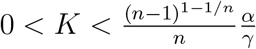.

*Proof*. Required steady states are the roots of

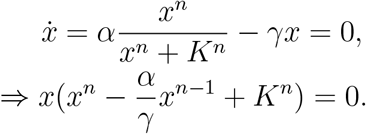

One steady state is at *x*_*o*_ = 0 and the other steady states are the roots of the polynomial 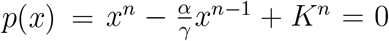. The roots of *p*(*x*) = 0 may be estimated via the method of root locus (Ogata, 2010), by rewriting it as,

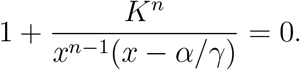

This is in a form from where the locus of roots as *K* is varied from 0 to ∞ may be plotted using standard methods, illustrated for *n* = 8 in Fig. 1. The root locus showed that there were two real positive roots when 0 *< K < K*_*c*_ and the remaining *n* − 2 roots were complex numbers. For *K > K*_*c*_, all roots were complex. To determine *K* = *K*_*c*_, we calculated the value of *x* where the root locus broke away from the positive real axis. The breakaway point *x*_*b*_ is a root of the polynomial *p*(*x*) and its derivative polynomial, denoted *p*^*′*^(*x*),

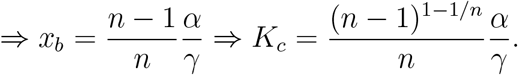

**Fig. 1:**
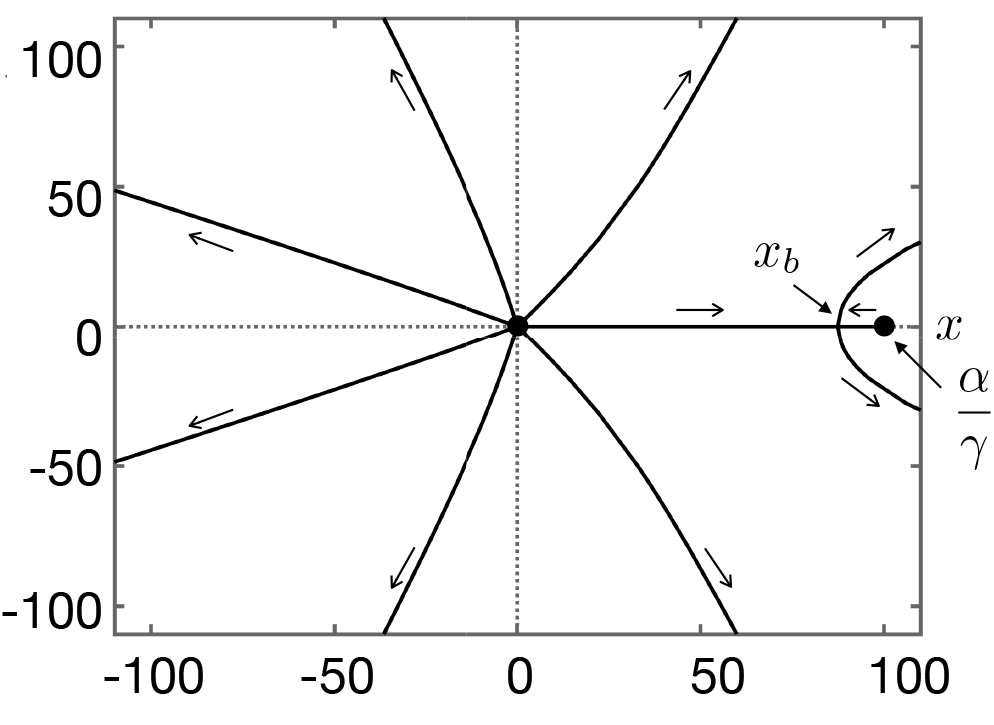
Root locus for *n* = 8, *x*_*b*_ represents the breakaway point.

The stability of the steady states automatically follows from geometric considerations. The above condition provides a simple check to determine whether bistable dynamics would persist under multi-parametric perturbations. As the nonlinearity is increased through an increase in *n*, the region of parameters exhibiting bistability increases (Fig. 2). Therefore, it agrees with the design principle that nonlinear positive feedback is needed for bistability. This proposition will help in designing synthetic bistable circuits that directly identify the parameters, ensuring bistability without the exact determination of the steady-state values.

**Fig. 2:**
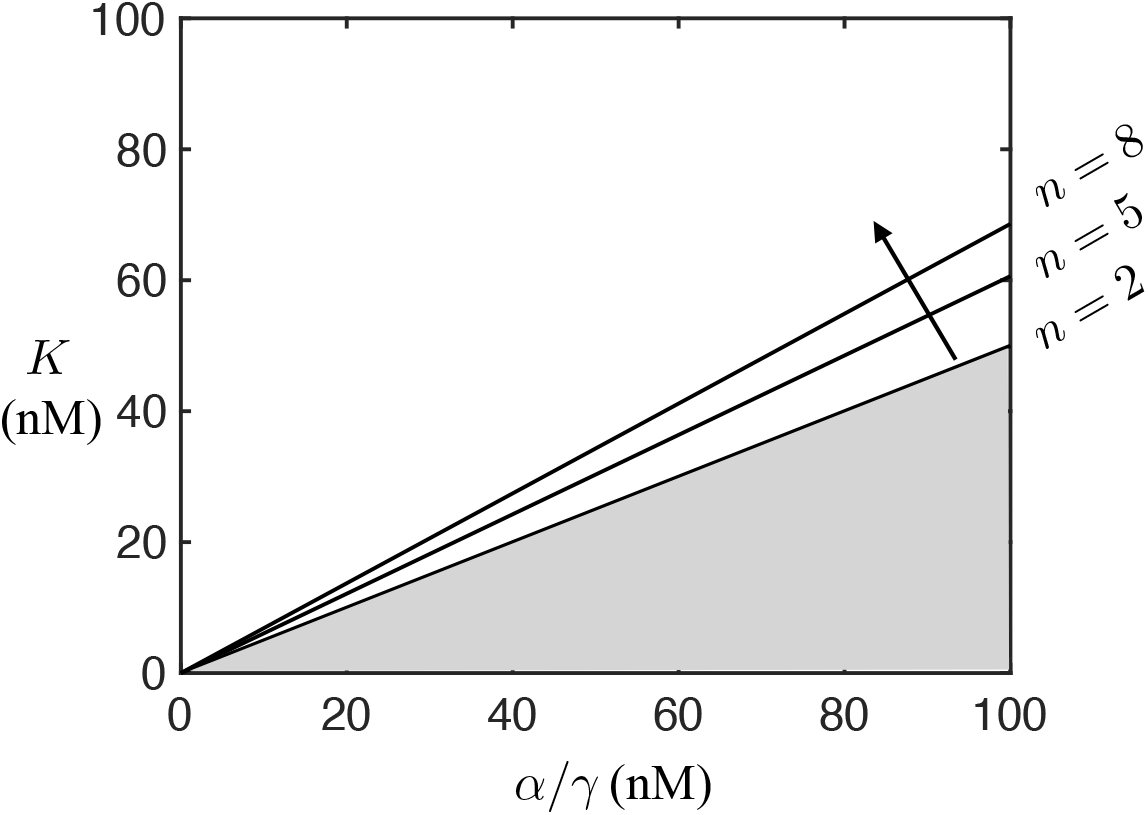
The shaded area represents the bistability region for *n* = 2. The region of bistability increases as the value of Hill coefficient *n* increases, as indicated.

### B. Quantitative Robustness Analysis

The exact steady state values would depend on the parameters values and perturbations to the parameters values would then propagate to the steady state values as well. The following proposition was obtained with the aim of getting a quantitative relation between the relative change in the unstable steady state and the relative change in the stable steady state in terms of the parametric perturbations.

#### Proposition 2.

*If δα, δγ, and δK are sufficiently small perturbations in the parameters α, γ, and K, respectively, then*,

1. 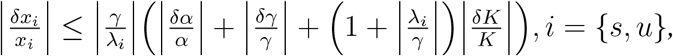
2. 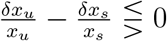 where 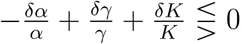 and

*Proof*. At *x* = *x*_*i*_, *i* = *{s, u*},

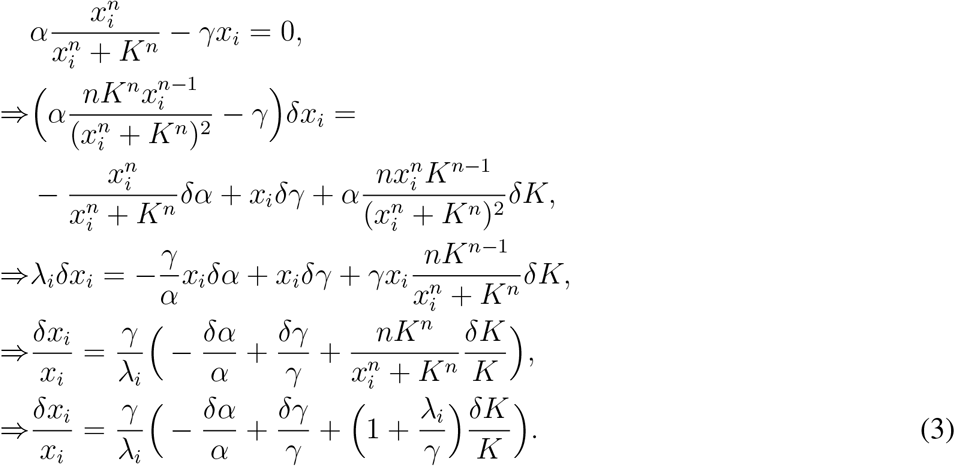

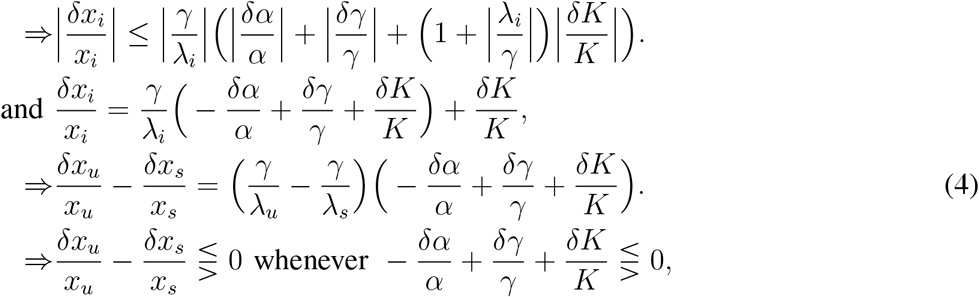

as *λ*_*u*_ *>* 0 *> λ*_*s*_. □

Equation (3) showed that the relative change of the steady state is proportional to a combination of the relative changes in the parameters, with the constant of proportionality being related to the eigenvalue at that steady state. The ratio of the relative change in the unstable steady state to the relative change in the stable steady state indicates how much more sensitive the unstable steady state is to parametric perturbations than the stable state. A larger magnitude of the eigenvalue at the unstable state indicates that this unstable steady state is generally more responsive to parameter perturbations compared to stable states. This explains why the bistable systems often have more delicate switching thresholds. For completeness, we listed in Table I the values of the stable steady state *x*_*s*_, the unstable steady state *x*_*u*_ and their eigenvalues for *n* = 2 and *n* → ∞. For *n* = 2, the steady-states given can be explicitly calculated using (1), and the respective eigenvalues can be derived from (2). However, for large *n* the hill function 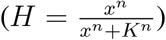 can be approximated as a step at *x* = *K*. We can write (1) as

**TABLE I:**
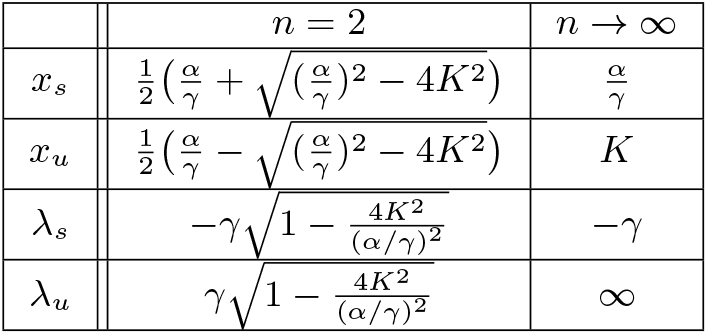
Steady states and eigenvalues.

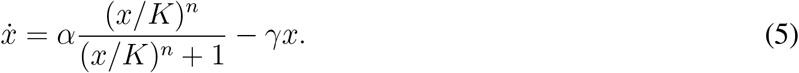

**Case 1:** For *x < K* implies (*x/K*)^*n*^ *<* 1. As *n* → ∞ implies (*x/K*)^*n*^ → 0, *H* → 0 such as 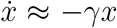 the steady-state *x*_*o*_ → 0 with eigenvalue as the first derivative of the approximated function *λ*_0_ → −*γx*,.

**Case 2:** For *x > K* implies (*x/K*)^*n*^ *>* 1. As *n* → ∞ implies (*x/K*)^*n*^ → ∞, *H* → 1 such as 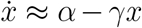, the steady-state *x*_*s*_ → *α/γ* with eigenvalue as the first derivative of the approximated function *λ*_*s*_ → − *γ*.

**Case 3:** *x* = *K* is the threshold state, which is the switching point of the Hill function (*H* → 1*/*2) for *n* → ∞, implies *x*_*u*_ → *K*, with the eigenvalue from (2) being 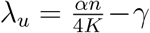, such as *n* → ∞, *λ*_*u*_ → ∞. The next proposition highlighted the effect of multi-parametric perturbations on the dynamics of bistability.

#### Proposition 3.

*The deviation in trajectories around the unstable state due to a parametric perturbation satisfies* 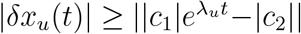 *where* 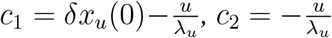, *and* 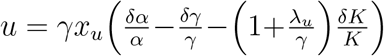.

*Proof*. The linearized dynamics around the unstable steady state in response to a multi-parametric parametric perturbation is of the form,

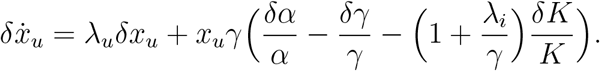

As 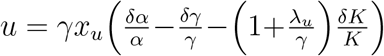

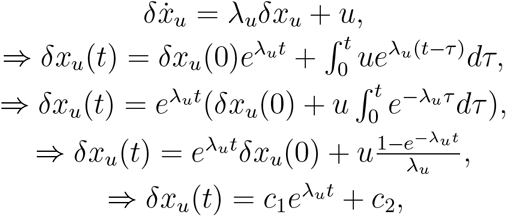

where 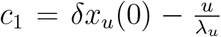 and 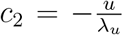. As long as the initial deviation is not exactly matched to the parametric perturbation (*c*_1_*≠* 0),

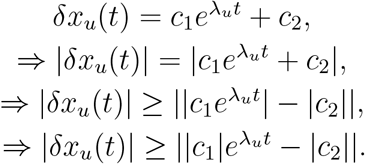

The unstable steady state is important for the transition from one stable steady state to the other stable steady state. Proposition (3) showed that in the neighbourhood of the unstable steady state, the multi-parametric perturbations cause a large change in the trajectories. This could have a significant impact in the overall transition from one stable state to another stable state. This sensitivity is directly related to the magnitude of the eigenvalue at the unstable steady state, with a large magnitude of the eigenvalue resulting in larger deviations faster. This shows an important design tradeoff, as increasing the nonlinearity increases the bistability region but makes the switching dynamics more susceptible to changes in parameters.

### C. Rigorously Bounded Deviations

The following proposition rigorously examines the extent to which the system’s steady states may vary when the parameters are allowed to vary over finite, bounded ranges. Relevant background on the interval analysis can be found in (Chorasiya et al., 2023).

#### Proposition 4.

*For the bistable positive feedback circuit* (1),

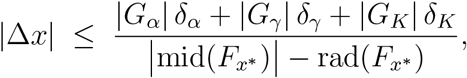

*where the nominal parameter gains are* 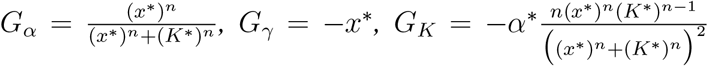, *and* 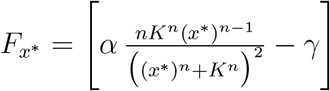 *is an interval extension of the Jacobian* 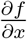 *at steady state x*^∗^.

*Proof*. Let *U* = [*α, γ, K*] be an interval vector of uncertain parameters with nominal value *u*^∗^ = (*α*^∗^, *γ*^∗^, *K*^∗^) and symmetric perturbation

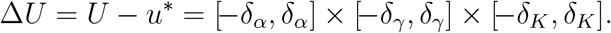

Let *x*^∗^ denote a steady state of Eq. (1). Furthermore, let *F*_*x*_∗ and *F*_*u*_∗ represent the interval extensions of the Jacobian matrices 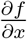 and 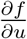 evaluated at (*x*^∗^, *u*^∗^), respectively.

Every admissible steady-state deviation Δ*x* from any multi-parametric perturbation Δ*u* ∈ Δ*U* lies in the interval solution set

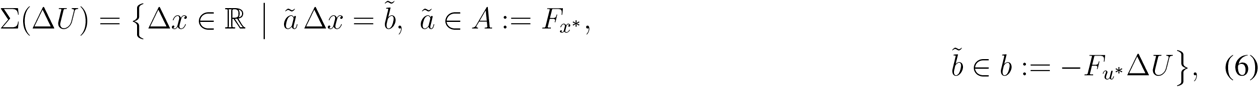

Consequently, every admissible Δ*x* satisfies the Oettli-Prager inequality (Chorasiya et al., 2023; Oettli and Prager, 1964)

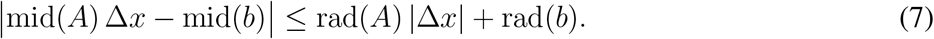

In this one-dimensional setting, *A* and *b* are interval scalars, so (7) becomes a scalar inequality in Δ*x*. Since the parameter intervals are symmetric around their nominal values, the forcing interval *b* = − *F*_*u*_∗ Δ*U* is centered at the origin, mid(*b*) = 0. This implies,

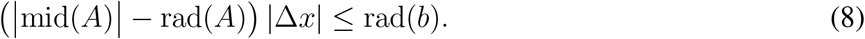

If mid(*A*) *>* rad(*A*) (equivalently, if *λ≠* 0),

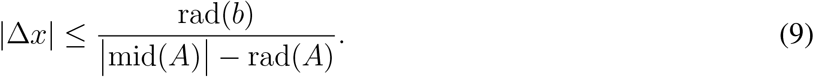

For the bistable positive feedback model (1), this results

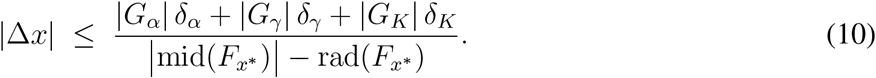

Thus, the interval in (10) offers a rigorous and fully explicit bound on the steady-state deviations, expressed in terms of the multi-parametric uncertainties (*δ*_*α*_, *δ*_*γ*_, *δ*_*K*_) and the Hill coefficient *n*.

The key implication of this approach is the capability to numerically and rigorously quantify the bounds of steady-state deviations in the presence of parametric uncertainties. In the numerical results presented in Table II, we employ interval Newton methods (Chorasiya et al., 2023) to obtain rigorous enclosures for *x*_*s*_ and *x*_*u*_, and to systematically quantify the dependence of these equilibria on increasing values of Δ*U*.

**TABLE II:** Rigorous bounds on steady states and relative variations for different levels of parameter uncertainty. Nominal parameter values: *α* = 100, *γ* = 1, *K* = 10.

| $\Delta U$ | $X_s$ | $\Delta x_s/x_s$ | $X_u$ | $\Delta x_u/x_u$ |
| --- | --- | --- | --- | --- |
| 0.1% | [99.80, 100.20] | [-0.002, 0.002] | [7.26, 7.29] | [-0.0015, 0.0015] |
| 1% | [98.02, 102.02] | [-0.02, 0.02] | [7.17, 7.38] | [-0.015, 0.015] |
| 2% | [96.08, 104.08] | [-0.04, 0.04] | [7.06, 7.49] | [-0.03, 0.03] |
| 5% | [90.47, 110.53] | [-0.1, 0.1] | [6.75, 7.82] | [-0.073, 0.073] |
| 10% | [81.81, 122.22] | [-0.2, 0.2] | [6.23, 8.39] | [-0.15, 0.15] |

Deriving the analytical expressions of steady-state values for *n* = 3, 4, 5, … would be challenging and algebraically complex. From these calculations, we noted that for *n* = 2, |*λ*_*s*_| = |*λ*_*u*_| and for *n* → ∞, |*λ*_*s*_ | ≪ |*λ*_*u*_|. As the magnitude of these eigenvalues have a significant impact on the relative change in the steady states and the sensitivity of the bistable dynamics, we computed these for other values of *n >* 2. We found that the unstable steady state always had an eigenvalue magnitude greater than or equal to that of the stable steady state (Fig. 3). These simulations imply the corollary |*λ*_*s*_| *<* |*λ*_*u*_| for Hill coefficients *n* = 3, 4, 5, 8, 10, 20, 50, consistent with Table I: as *n* increases, |*λ*_*s*_| converges to a finite limit *γ*, while |*λ*_*u*_| diverges to infinity. We used a rigorous numerical construction using interval analysis to prove statements about the magnitude of *λ*_*s*_ and *λ*_*u*_. The interval Newton algorithm was employed to rigorously determine quantitative bounds on the steady states. Following the determination of the steady states, (2) was used to compute bounded eigenvalues.

**Fig. 3:**
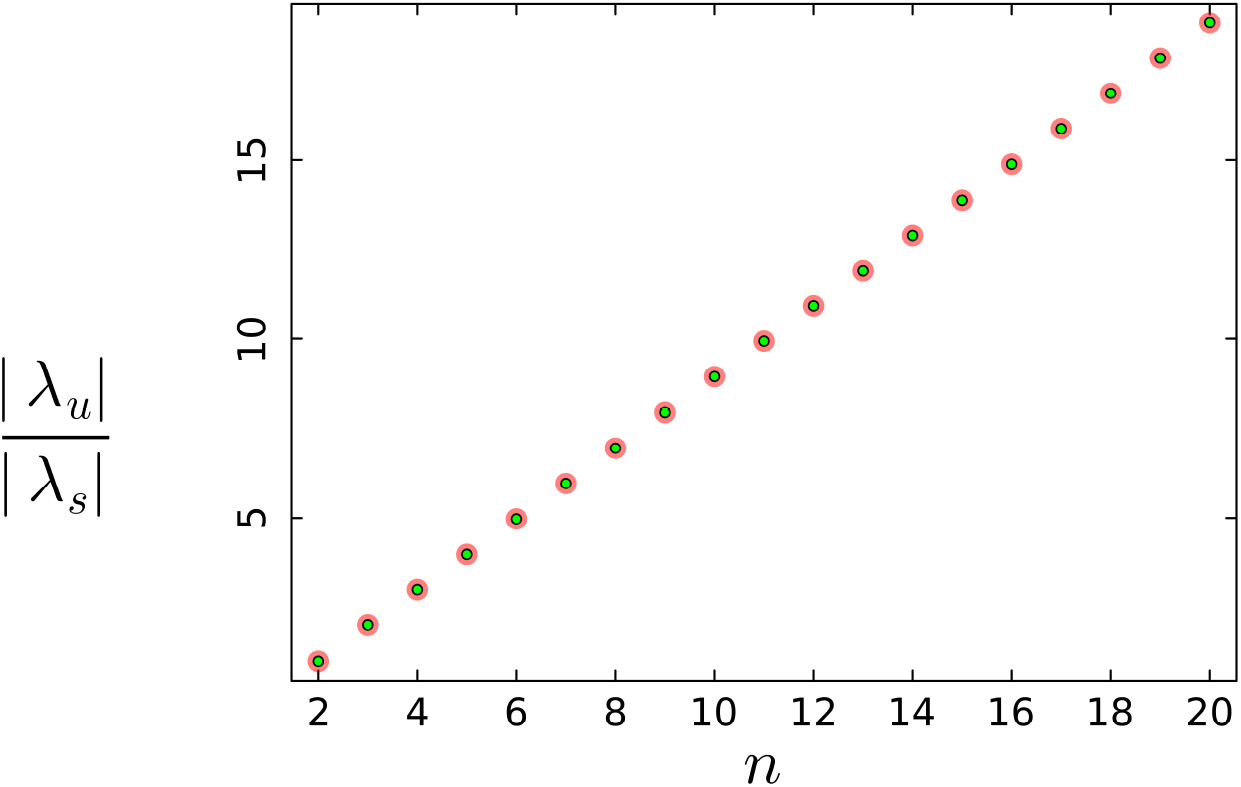
Relative magnitude of the eigenvalues for different *n*. Parameters: *α* = 100 nM/hr, *γ* = 1/hr, *K* = 1 nM. Red and lime colour markers indicate interval-valued and real-valued calculations, respectively.

#### Corollary 1.

*For α* = 100, *γ* = 1, *K* = 1, *and n* = 3, 4, 5, 8, 10, 20, 50, |*λ*_*s*_| *<* |*λ*_*u*_|.

*Proof*. The interval Newton algorithm (Chorasiya et al., 2023) was employed to rigorously determine the quantitative bounds of the steady states. Furthermore, (2) was used to compute bounded eigenvalues; see Table III. It is evident that, for all values of *n*, the inequality |*λ*_*s*_| *<* |*λ*_*u*_| holds.

**TABLE III:**
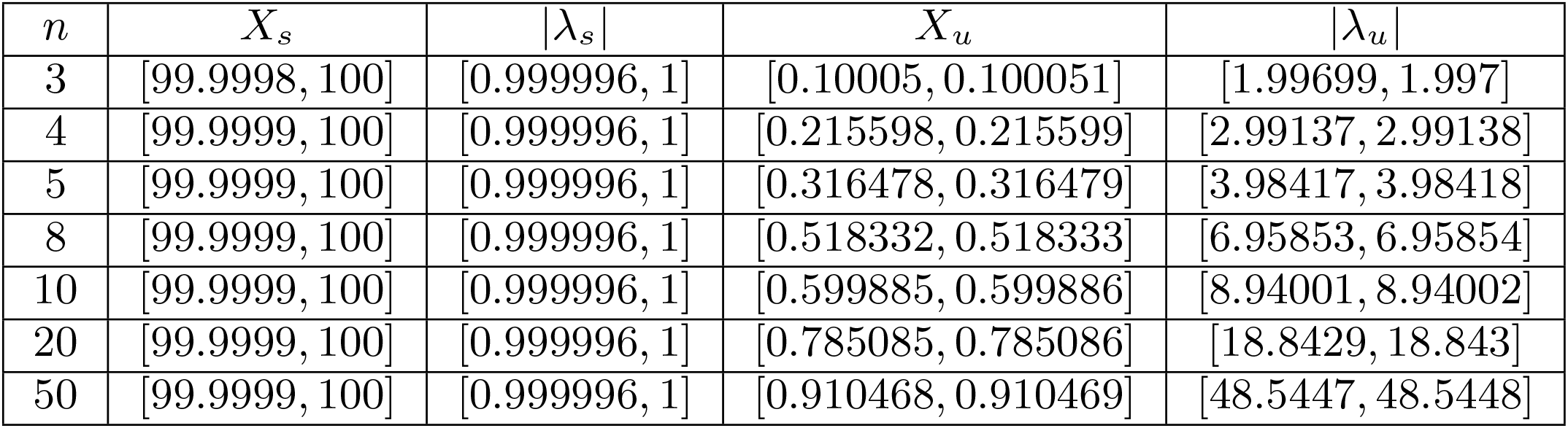
Bounded steady states and their eigenvalues.

This comparison shows that while the final stable cellular fates have slow dynamics and are robust to changes in *n*, the separatrix between them acts as a highly sensitive switch. Its dynamics become faster and more unstable, with a higher positive *λ*_*u*_, as cooperativity increases.

Corollary (1), together with Figure 2 and 3, supports the conclusion that there is an inherent trade-off: as the nonlinearity (*n*) increases, the region of parameter space exhibiting bistability expands, but the resulting bistable dynamics become correspondingly more sensitive (*λ*_*u*_ increases). Nonlinear positive feedback is the standard design principle for bistability, but it inherently creates sensitivity focused at the threshold state. Paradoxically, the same design choices that stabilise the existence of bistability also exponentially amplify this vulnerability.

The interval-analysis-based method provides an effective approach to examine the impact of parametric uncertainty on the complete time trajectory of the system. To this end, we represent parametric uncertainty in the nonlinear differential equations using interval-valued parameters. We then compute the system trajectories for the specified parameter intervals and intervals of initial conditions using a Taylor-model-based reachability analysis method (Chen, 2015). All computations are performed in Julia 1.10.5 using the package ReachabilityAnalysis v0.27.0.

The simulation results are shown in Fig. 4. The figure depicts the state trajectories of a bistable positive feedback system subject to multi-parametric uncertainty, evaluated for different values of the Hill coefficient *n*. The initial condition is specified as *X*_0_ ∈ [4.5, 4.5], with uncertain parameter intervals given by *α* ∈ [100, 101], *γ* ∈ [1, 1.01], and *K* ∈ [10, 10.1]. For each *n* (e.g., *n* = 2, 3, 4), the figure plots the bounded time trajectories from the initial condition over the given parameter ranges.

**Fig. 4:**
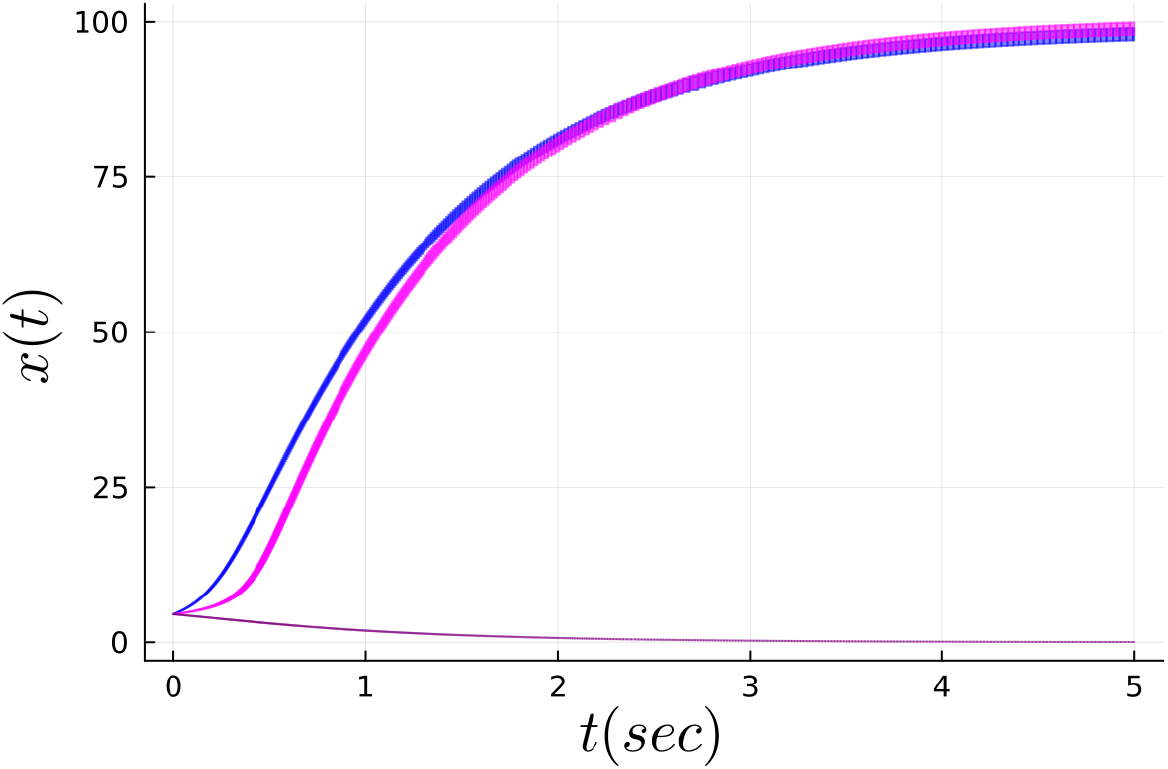
For the parameter intervals *α* ∈ [100, 101] nM/hr, *γ* ∈ [1, 1.01] hr^−1^, *K* ∈ [10, 10.1] nM, and initial condition *X*_0_ = [4.5, 4.5], the corresponding bounded-time trajectories are displayed for different values of *n*. Using the same initial condition *X*_0_ and identical parameter intervals, the trajectories depicted in blue and magenta exhibit distinct transient slopes for *n* = 2 and *n* = 3, respectively. However, the trajectory shown in purple, corresponding to *n* = 4, converges to zero. This indicates that the system’s sensitivity during the transient phase depends strongly on the Hill coefficient *n*.

The trajectories from *X*_0_ change qualitatively with *n*. Using the same initial condition, *X*_0_, and identical parameter ranges, the trajectories shown in blue and magenta display noticeably different transient slopes for *n* = 2 and *n* = 3, respectively. As *n* increases, the slope observed during the transient phase becomes steeper. In contrast, the purple trajectory, corresponding to *n* = 4, decays to zero. Together, these observations suggest that the system’s transient behaviour is highly sensitive to the value of the Hill coefficient *n*. Therefore, the system is bistable, with two stable steady states separated by an unstable one. The unstable steady state (*x*_*u*_) represents the critical threshold that the system must cross to transition between cellular states. The outcome from a given initial condition is highly sensitive to *n*. Because varying *n* can change whether trajectories stay bounded or diverge and whether they reach the lower or higher stable branch, the robustness of bistability depends strongly on the Hill coefficient, underscoring the key role of cooperativity.

## III. COMPUTATIONS

We computed the steady states and eigenvalues of (1) to check if they agree with the predictions summarised in the propositions above. The steady states were computed using the rlocus command in MATLAB, and the dynamics were simulated using ode23s. Fig. 5 showed that the relation between the relative change in the unstable steady state and the relative change in the stable steady state was consistent with expectation in terms of the combination of relative changes in the parameters. Fig. 6 showed the almost exponential divergence of the trajectories in the linearized dynamics approximated the difference in the nonlinear model quite well. Fig. 7 showed the impact of this exponential divergence in terms of the actual difference in the dynamics of the perturbed and unperturbed trajectories with both deterministic and stochastic simulation. The stochastic simulations were done using Gillespie’s algorithm (Vecchio and Murray, 2014).

**Fig. 5:**
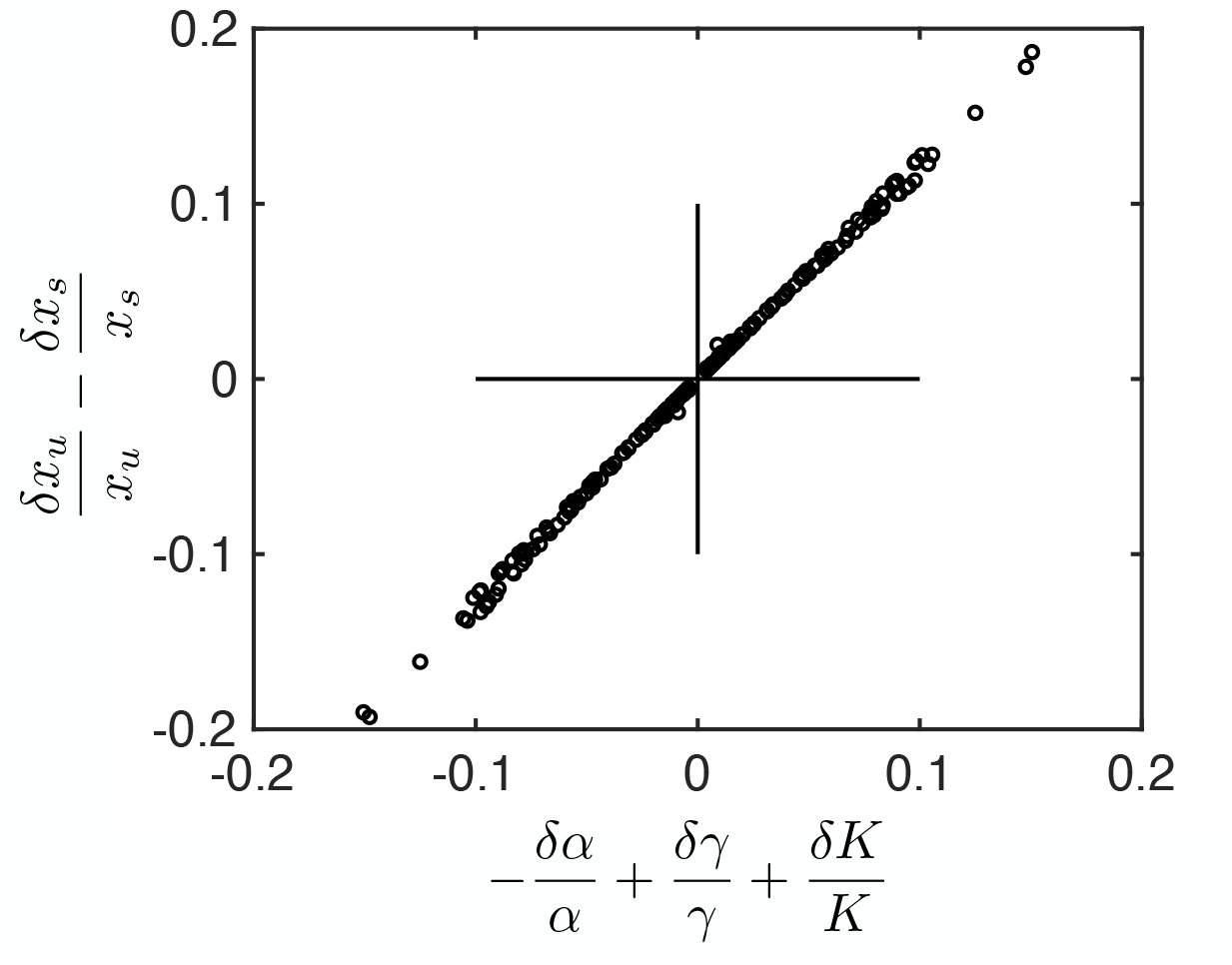
Relative change in steady states due to parametric perturbation. Each circle represents the computation from one parameter set. Default parameters: *α* = 100 nM/hr, *γ* = 1/hr, *K* = 1 nM. The parameter space was sampled by randomly multiplying each of the above parameters by a uniformly randomly chosen number between [10^−1^, 10^1^]. Only those parameter sets that exhibited bistability were considered. For each such parameter set, the parameters were perturbed by *h* = 0.01 or *h* = − 0.01. The solid lines represent guidelines that identify the different quadrants.

**Fig. 6:**
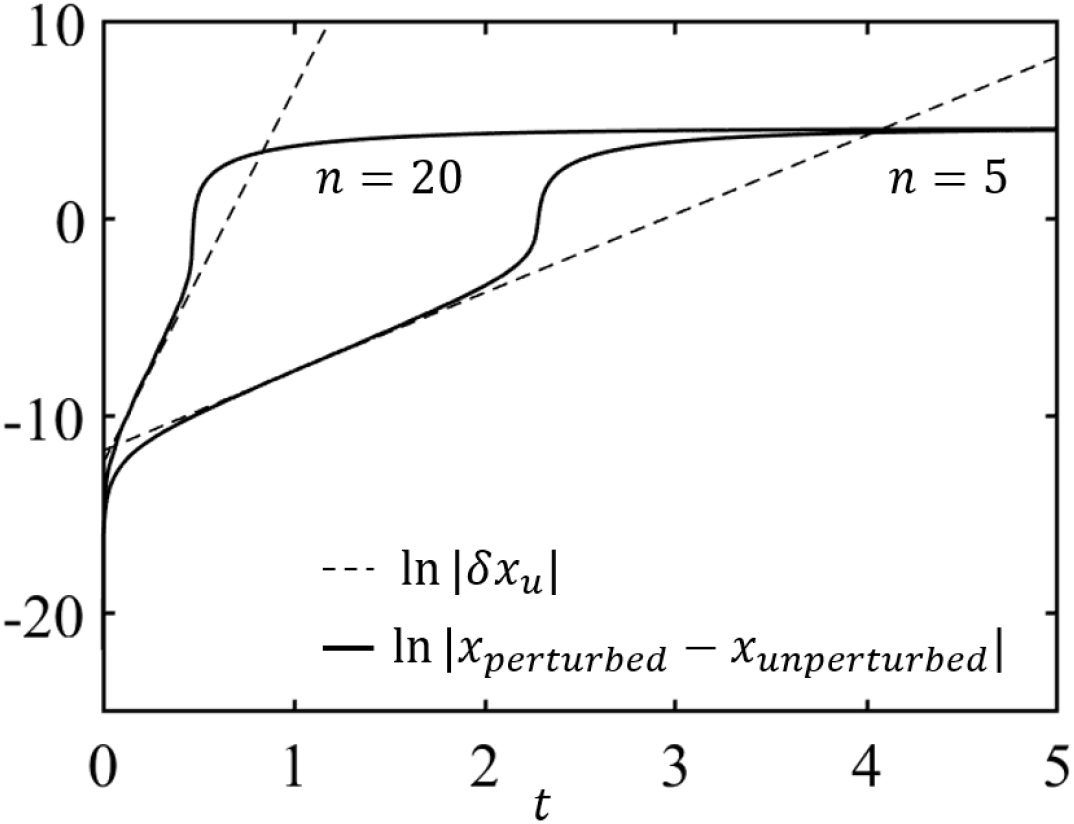
Exponential divergence of trajectories due to perturbation. Dashed line represents the absolute value of the trajectory obtained from the linearized dynamics. The initial condition was taken as the difference between the numerically computed values of the unstable steady state (*x*_*u*_) of the perturbed and unperturbed system. Solid line represents the absolute value of the difference of the trajectories obtained from the perturbed and unperturbed systems using the full nonlinear dynamics. The initial conditions for both the full nonlinear dynamics simulations was taken as the numerically computed value of the unstable steady state (*x*_*u*_) of the unperturbed system. Parameters: *α* = 100 nM/hr, *γ* = 1/hr, *K* = 1 nM, *n* = 5 and *n* = 20. The perturbation was to increase the value of *α* by *h* = 0.01.

**Fig. 7:**
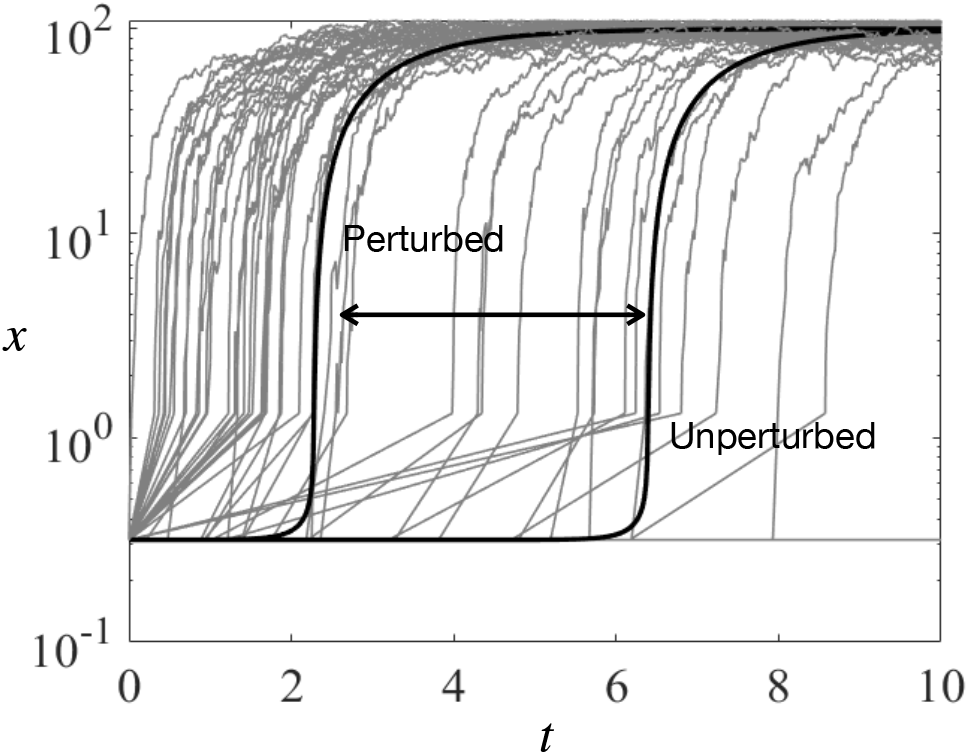
Difference between the perturbed and unperturbed trajectories can be significant. Solid black lines represent the trajectories from the full nonlinear dynamics of the unperturbed and perturbed systems, as indicated. Grey lines represent stochastically simulated trajectories (*m* = 50) that also show the variations with the same parameters as used in deterministic trajectories. The initial condition was taken as the numerically computed value of the unstable steady state (*x*_*u*_) of the unperturbed system. Parameters: *α* = 100 nM/hr, *γ* = 1/hr, *K* = 1 nM, *n* = 5. The perturbation was to increase the value of *α* by *h* = 0.01.

## IV. DISCUSSION

We investigated the quantitative aspects of robustness to bistable dynamics to multi-parametric perturbations in a benchmark biomolecular positive feedback model. Using the method of root locus, we obtained a necessary and sufficient condition for the existence of bistability in terms of the model parameters. We obtained a quantitative relation between the relative change in the stable steady state and that in the unstable steady state in terms of the relative perturbation in the parameters. We showed that the deviation in the trajectories near the unstable steady state due to parametric perturbations increased almost exponentially, corresponding to the eigenvalue, but eventually bounded by the nonlinear dynamics. Using interval integration of nonlinear differential equation we enclosed the complete, transient and steady state response of the system. These results were illustrated using numerical simulations. Using validated numerical construction, we showed that the magnitude of the eigenvalue of the unstable steady state is always greater than that of the stable steady state. In particular, the behaviour of the system near the unstable steady state is highly sensitive to changes in multiple parameters, effectively acting as a delicate pivot that determines how the system switches between states. This reveals a fundamental design trade-off: increasing the degree of nonlinearity expands the range of parameter values over which bistability can be maintained, but at the same time it makes the threshold state more unstable, and thus makes the switching dynamics exponentially more susceptible to perturbations.

